# Isolation and characterization of persisters of the pathogenic microorganism *Staphylococcus aureus*

**DOI:** 10.1101/2023.09.19.558453

**Authors:** Shiqi Liu, Paul Laman, Sean Jensen, Nicole N. van der Wel, Gertjan Kramer, Sebastian A. J. Zaat, Stanley Brul

**Affiliations:** Department of Molecular Biology and Microbial Food Safety, Swammerdam Institute for Life Sciences, University of Amsterdam, Science Park 904, 1098 XH Amsterdam, The Netherlands; Department of Medical Biology, Electron Microscopy Center Amsterdam, Academic Medical Center, University of Amsterdam, Meibergdreef 9, 1105 AZ, Amsterdam, the Netherlands; Department of Mass Spectrometry of Biomolecules, Swammerdam Institute for Life Sciences, University of Amsterdam, Science Park 904, 1098 XH Amsterdam, The Netherlands; Department of Medical Microbiology, Centre for Infection and Immunity Amsterdam (CINIMA), Academic Medical Centre, University of Amsterdam, Meibergdreef 9, 1105 AZ Amsterdam, The Netherlands

**Author notes:** These authors contributed equally to this work (Sebastian A. J. Zaat, Stanley Brul).

**Keywords:** antibiotic persistence, *Staphylococcus aureus*, isolation, proteomics, characteristics

## Abstract

*Staphylococcus aureus* is a notorious pathogen responsible for significant morbidity and mortality in both human society and animal husbandry. The presence of *S. aureus* persisters is also one of the leading causes of recurrent and chronic diseases. Persisters are a subset of growth-arrested bacteria within a susceptible bacterial population that are able to tolerate antibiotic treatment and resuscitate after stress removal. Consequently, investigating their formation and characteristics is of crucial importance to provide mechanism-based options for their eradication. However, one challenge in mechanistic research on persisters is the enrichment of pure persisters. In this work, we validated a proposed method to isolate persisters from vancomycin and enrofloxacin generated persistent populations. With this, we analyzed the proteome profile of pure persisters and revealed the distinct mechanisms associated with vancomycin and enrofloxacin induced persisters. Furthermore, morphological and metabolic characterizations were performed, indicating further differences between these two persister populations. Finally, we assessed the effect of ATP repression, protein synthesis inhibition and reactive oxygen species (ROS) level on persister formation. In conclusion, this work provides a comprehensive understanding of *S. aureus* vancomycin and enrofloxacin induced persisters at the molecular, single cell and population levels, facilitating a better understanding of persisters and the development of effective strategies to combat them.

## Introduction

*Staphylococcus aureus*, a well-known and prevalent pathogen, is responsible for a variety of infections of e.g., human skin, joints, and bloodstream as well as infections occurring in animal husbandry. This results in considerable morbidity and mortality [1]. According to a recent review, almost 1.2 million individuals in the United States alone suffered from *S. aureus* bloodstream infections in 2017, with a fatality rate of one-sixth among those patients [2]. Unfortunately, *S. aureus* is also one of the leading pathogens causing chronic infections, primarily due to the presence of persisters. The term “persister” was coined by Joseph W. Bigger in 1944 [3] when he found that in a penicillin-treated *S. aureus* population, a small proportion of cells prevented to attain sterility, without however becoming resistant. Approximately a decade later, the correlation between persisters and prolonged antimicrobial therapy began to draw attention in both animal models and human patients [4,5]. Until now, persisters have been found in almost all tested bacteria and, as stated, are regarded as the main causative agent of chronic diseases and recurrent contamination [6].

For a long time, persisters were defined as a dormant phenotype with negligible metabolic activity and the dormancy itself was claimed as the primary reason of stress tolerance. However, increasing evidence emerged that growth arrested persisters do exhibit a certain level of metabolic activity [7]. Our previous work demonstrated that in a stable persistent population, despite being growth arrested, *S. aureus* cells were able to modulate various metabolic pathways at different time points to combat different stressors, resulting in distinct molecular features in each persistent population [8]. This stress related coping mechanism can be considered as a double-edged sword. On the bright side, specific stress responses towards one particular drug can potentially render cells more vulnerable to another kind of antibiotic due to the energy demands for such stress coping mechanism. This may make combined therapy a promising method to act synergistically in the reduction of persistence levels. However, on the other hand, this phenomenon limits the full development and utilization of antimicrobial agents that may rely for their activity on a certain level of cell growth. Therefore, it is crucial to uncover the molecular and cellular features of persisters generated from cells exposed to different stress conditions such that knowledge-based strategies for their eradication can be formulated.

Given that persisters contain a similar genome composition compared to control cells, proteomics could work as a powerful tool to elucidate mechanisms behind this stress tolerant phenotype and its formation. However, due to the low frequency of persisters and their transient nature, the utilization of proteomics has encountered numerous challenges as reviewed in [9]. One challenge in persister research is the enrichment of persisters. In the stated previous work, we treated *S. aureus* populations with high levels of antibiotics for 24 hours where most cells were killed within 4 hours of incubation, subsequently gradually fragmented, and then largely removed during routine washing steps. This led to final populations where the majority (>75 %) of cells were persisters, making it possible to analyze the persisters generation and some of their features without extra enrichment steps. However, this level of persister generation in antibiotic exposed populations is not universally applicable to any bacterial population or strategies to generate persisters. For example, in our previous work analyzing *Bacillus subtilis* persisters generated by different antibiotics, we observed that in the final washed population, the percentage of persisters ranged from 25 % to 90 % [10]. Therefore, sorting techniques such as Fluorescence-activated cell sorting (FACS) or Magnetic-activated cell sorting (MACS) combined with selected probes have been conducted to separate persisters [11,12].

A commonly employed sorting approach is to use the live/dead dyes SYTO9 and propidium iodide (PI), respectively, and consider the SYTO9 positive cells as persisters [11,13]. Another method is to select alive cells with lower or non-detectable metabolic activity compared to vegetative cells of the same [14]. However, it is important to note that these selected populations contain not only persisters, but likely also VBNC cells, which are also growth arrested and antibiotic tolerant. For instance, in ampicillin treated *Escherichia coli* cultures, more VBNC cells were found than persisters [15]. Ayrapetyan et al. [16] proposed a dormancy continuum model that encompasses persisters and more dormant VBNC cells because several phenotypic characteristics and molecular mechanisms are associated with both populations. However, the fact that persisters can easily resume growth in standard culture media after stress removal while VBNC cells have lost their ability to be cultured, demonstrates the distinction of these two populations. The ability to be resuscitated has been utilized to separate persisters from VBNC cells. For example, single-cell imaging and microfluidics allow the detection of culturable persisters and sleeping VBNC cell for physiological study at the single cell level [17–19]. Alternatively, inducible fluorescent signals from reporter proteins can be used to monitor bacterial division and thereby the presence of persisters and/or VBNC cells[20–22]. However, these methods alone are insufficient for isolating persisters for further analysis. Hence, finding a specific marker or probe that target persister-specific characteristics is of great importance. To that end the availability of purified persisters is a prerequisite.

In our previous study, we presented a persister isolation method with a commonly used model strain *Bacillus subtilis* where metabolism dependent dye 5-(and-6)-Carboxyfluorescein diacetate (CFDA) and PI detecting membrane permeabilization were used to label and isolate persisters [10]. Thereby, persisters can be analyzed at both single cell level and population level as purified cells. In the present work, we extend this method to the pathogenic microorganism *S. aureu*s. Upon generating and isolating persisters with this method, we performed proteomics of the isolated persister cells. This analysis revealed specific proteome features associated with different stress conditions, allowing us to compare and contrast the proteomic profiles of vancomycin and enrofloxacin persisters. Additionally, we further investigated the morphological and metabolic characteristics of persisters at both single cell and population levels. These data aided in the validation of various findings from the proteomics study. By combining proteomic, morphological, and metabolic investigations, we gained a comprehensive molecular physiological understanding of *S. aureus* vancomycin and enrofloxacin induced persisters. The findings showed that we are able to isolate *S. aureus* persisters and to understand their differential molecular response to 2 antibiotics, highly relevant to our knowledge of persister formation and survival mechanisms. The results potentially facilitate the development of effective treatments against persistent infections.

## Results

Previously, we analyzed the formation of *S. aureus* persisters through 24 hours of vancomycin or enrofloxacin exposure [8]. Although the modes of action of these two drugs are different, they generated similar levels of persisters with a final survival fraction of approximately 0.1 %. Distinctive molecular dynamics were observed for the individual biphasic kill curves.

### *S. aureus* persisters isolation and regrowth

In order to study pure persisters, we firstly evaluated the proposed CFDA-PI double staining method [10] with stable vancomycin and enrofloxacin induced *S. aureus* persistent populations. After 24 hours of antibiotic exposure, surviving cells were stained and inoculated in fresh Tryptic Soy Broth (TSB) at a ratio of 1:20 (stained culture: fresh medium). Double stained vegetative cells incubated in PBS were used as negative controls. CFDA intensities were detected by flow cytometry before (0h) and after 1.5h, 2h and 2.5h of incubation (Figure 1A). The proportions of CFDA^+^-PI^-^ cells in negative control and the 2 antibiotic treated groups at each time point were normalized to the corresponding 0h samples (Figure 1B). Altogether, compared with the non-changing CFDA signal in the negative control, the decrease of the CFDA signal in the CFDA^+^-PI^-^ population, where the change of CFDA intensity was deemed to be due to the quick regrowth of surviving cells after stress removal, was in agreement with the definition of persisters being antibiotic tolerant and able to quickly regrow after stress removal. Hence, for the following experiments, CFDA-PI double staining was used to isolate persisters from vancomycin and enrofloxacin treated samples, referring to them as van-P and enro-P, respectively. The growth curves of 10^5^ FACS-isolated persisters and vegetative cells were measured during 12h of incubation in TSB (Figure 1C). Compared with vegetative cells, persisters had a longer lag time but similar growth rates in the log phase. In addition, van-P had a slightly shorter lag time than enro-P.

**Figure 1.**
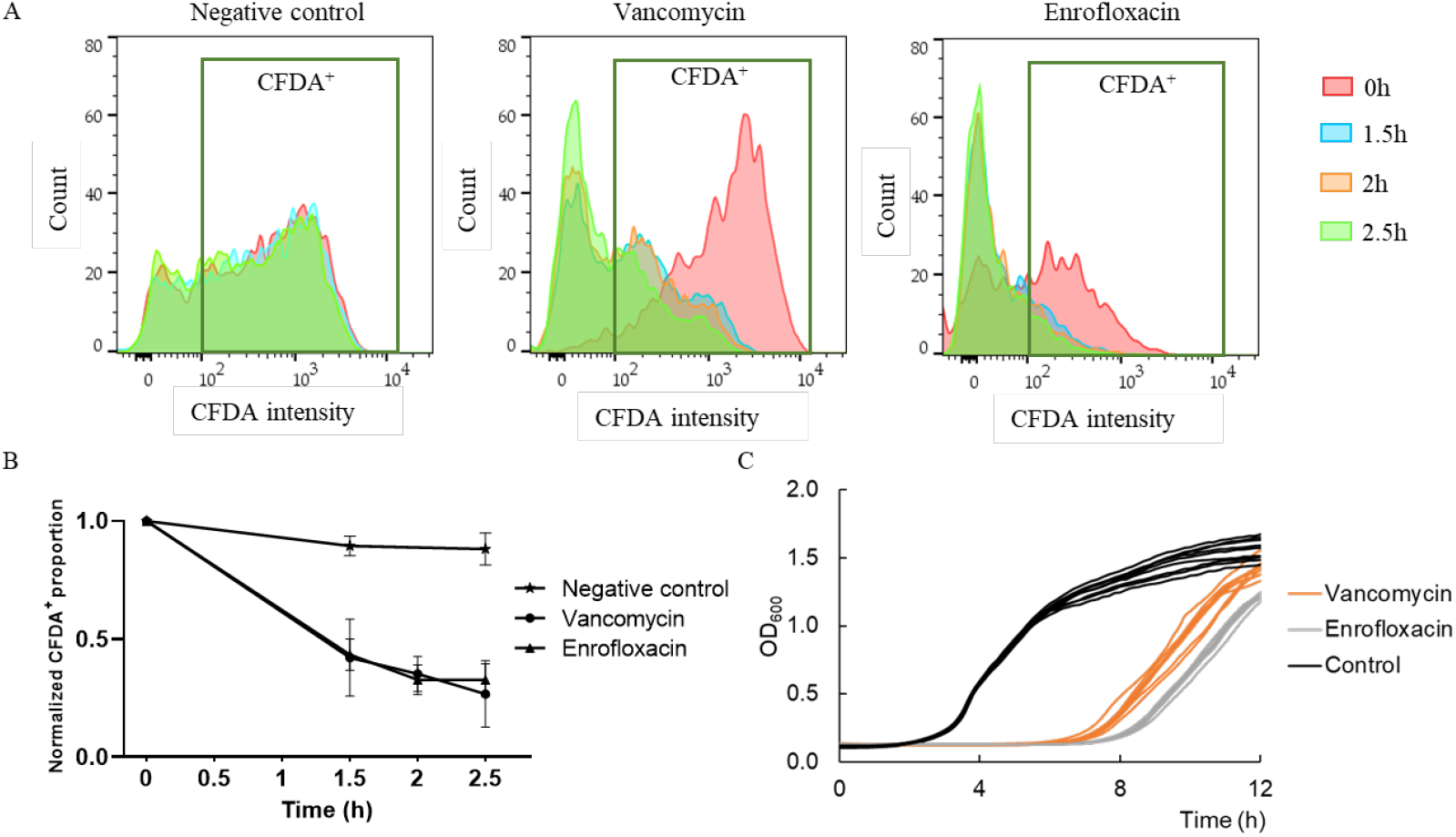
CFDA-PI double staining is able to label *S. aureus* persistent cells and identify the regrowth of persisters. **A**. After antibiotic exposure and CFDA-PI double staining, cultures were 1:20 inoculated into either PBS as negative control or fresh TSB liquid medium for cultivation. CFDA and PI intensity was measured at different time points by flow cytometry. In each sample, ten thousand events were analyzed. CFDA intensity higher than 10^2^ is considered as CFDA^+^. **B**. The proportions of CFDA^+^-PI^-^ cells were normalized against 0h control. In total, the constant CFDA signal in the negative control, and the decrease of the CFDA^+^-PI^-^ cells upon exposure of the population to TSB indicate that the change of CFDA intensity is due to regrowth of surviving cells after stress removal. This matches the definition of persisters. n≥2. **C**: Growth curve of Fluorescence-Activated Cell Sorting (FACS) isolated CFDA^+^-PI^-^ cells. 200 μl Fresh TSB medium containing 10^5^ isolated cells from vancomycin treated samples (orange), enrofloxacin treated samples (gray) and control vegetative cells samples (black) were incubated in 96-well plates for 12 hours at 37 ℃. OD_600_ values were measured every 10 minutes. N=9.

### Proteome profile of *S. aureus* persisters

To determine the proteome profile of van-P and enro-P cells, we utilized CFDA-PI double staining and FACS to isolate a pure persister population for quantitative proteomics. In total, 162 proteins were considered as valid out of 451 detected proteins. Among them, 109 proteins were identified from ctrl samples, 137 from van-P and 72 from enro-P. The coefficient of variation (CV) values (Figure 2A) indicates that the 3 biological replicates in each group had a positive correlation. To analyze the differences between persisters and vegetative cells, we displayed the data in a Venn diagram (Figure 2B) and applied a t-test (Figure 2 C&D) comparing control cells and persisters. In view of the limitation of cell number and sequencing depth, uniquely expressed and differentially increased proteins in persistent samples were both considered as factors that associated with persisters, while proteins only detected or higher expressed in control likely function as persistence repressors, or are dispensable for persister formation.

**Figure 2.**
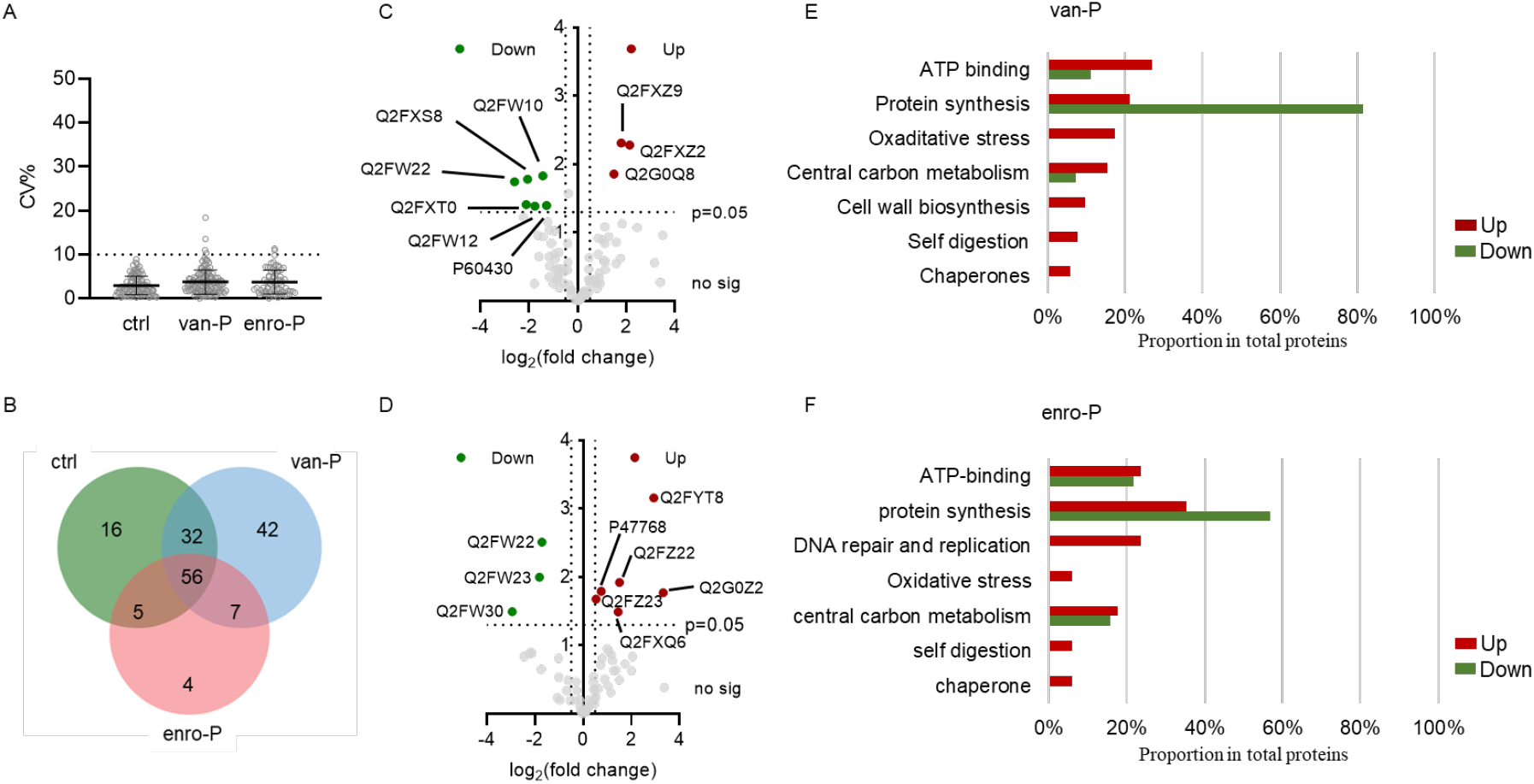
General comparison between proteomes of vancomycin induced persisters (van-P), enrofloxacin induced persisters (enro-P) and controls (ctrl). **A**. The coefficient of variation (CV) was calculated for proteins in each group. CV% <10% indicate that triplicates displayed a positive correlation. **B**. Venn diagram of proteins from each group. Volcano plots show the differentially expressed proteins in vancomycin persisters (**C**) and enrofloxacin persisters (**D**) compared with vegetative cells. Uniquely expressed proteins as well differentially expressed proteins in van-P (**E**) and enro-P (**F**) were grouped based on their functions. Red bars represent proteins that contribute to persister formation while green bars are proteins with lower expression in persisters. Single proteins may be included in more than one groups. The proportion refers to the ratio of the number of proteins in a certain category to the number of total proteins.

Compared to the 109 proteins found in the control group, the Venn diagram demonstrated that 49 proteins were exclusively expressed in van-P and 11 in enro-P. In addition, nine differentially expressed proteins in van-P (Figure 2C) and nine in enro-P (Figure 2D) were found. In total, vancomycin triggered the higher expression of 53 proteins and repressed 27 proteins (listed in Table 1), while in enro-P we observed 16 induced proteins and 51 decreased proteins (Table 2). Proteins with different functions were then classified and visualized in figure 2.3E and figure 2.3F, for van-P and enro-P, respectively.

**Table 1.**
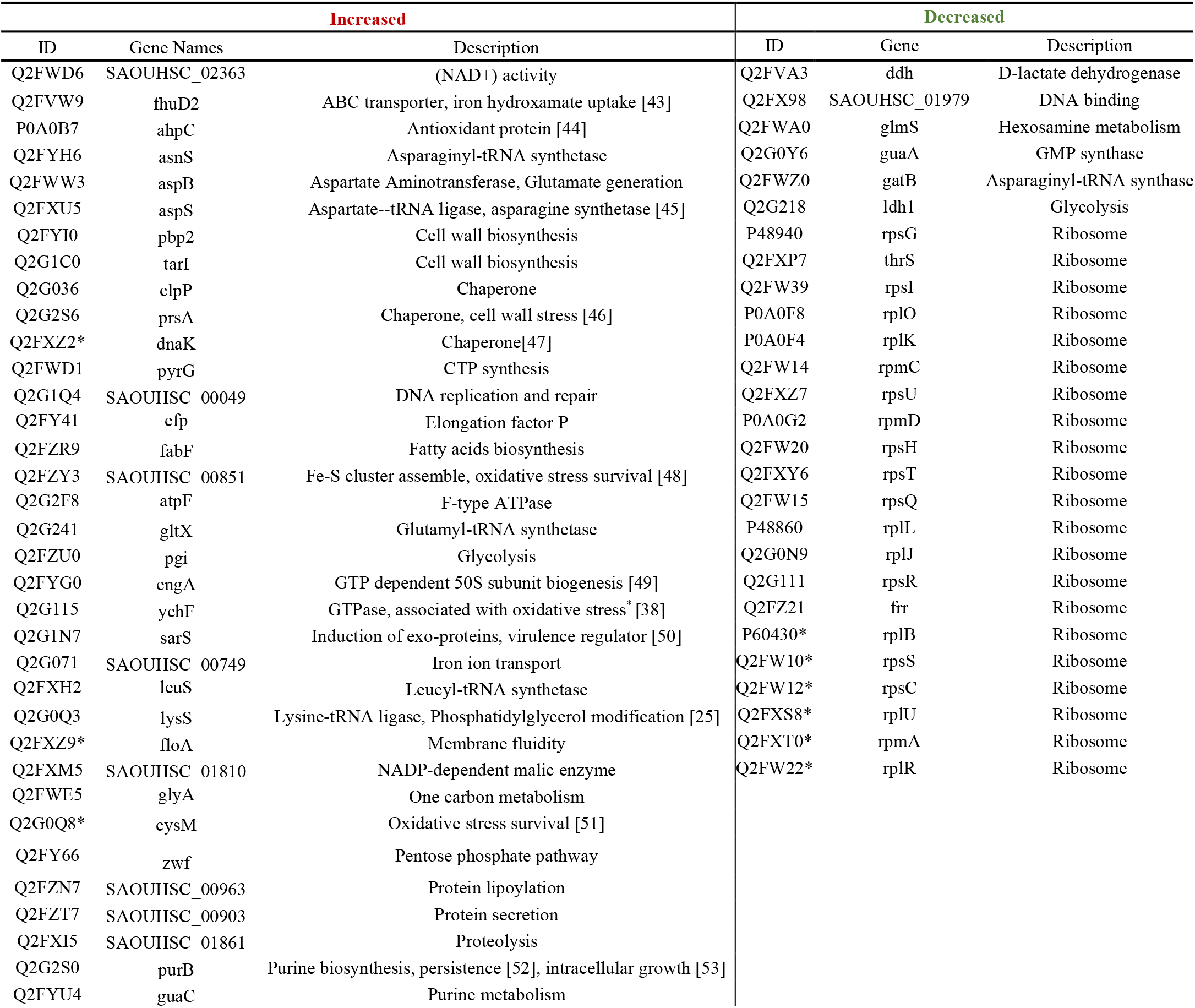
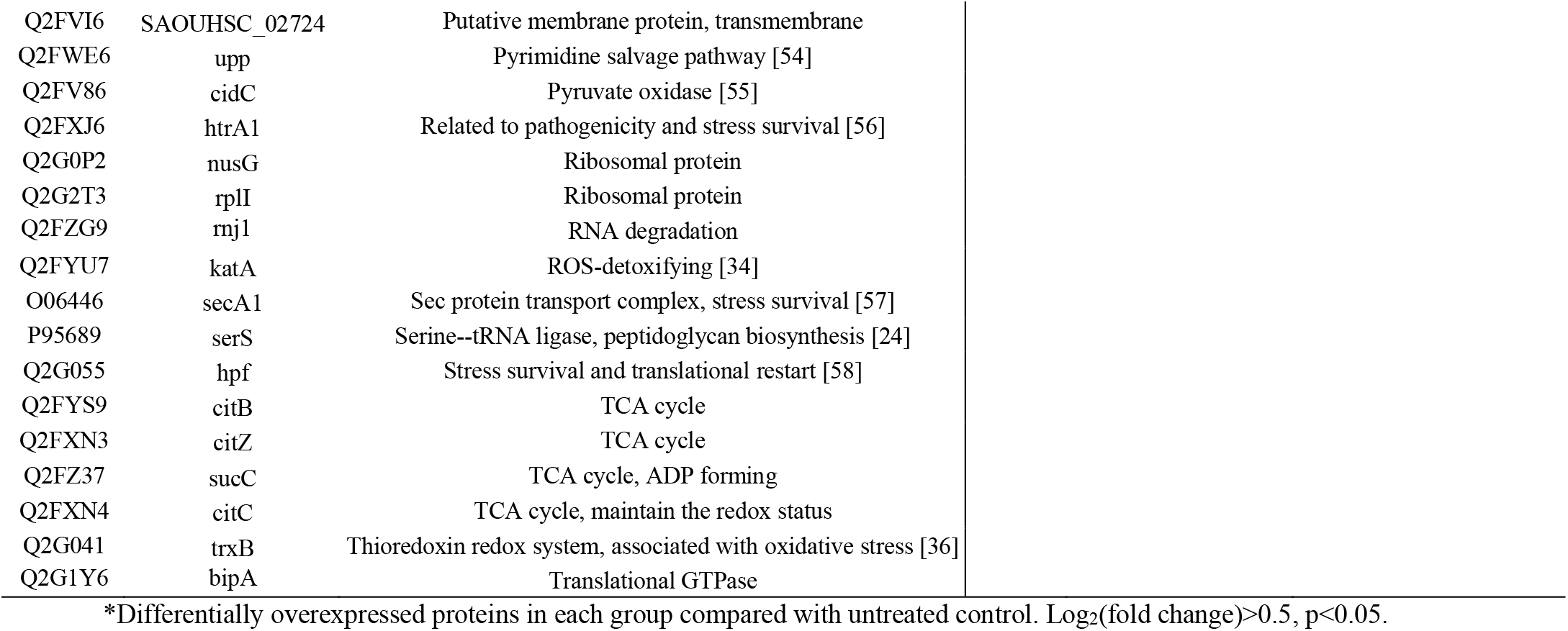
Increased or decreased proteins in vancomycin persisters.

**Table 2.**
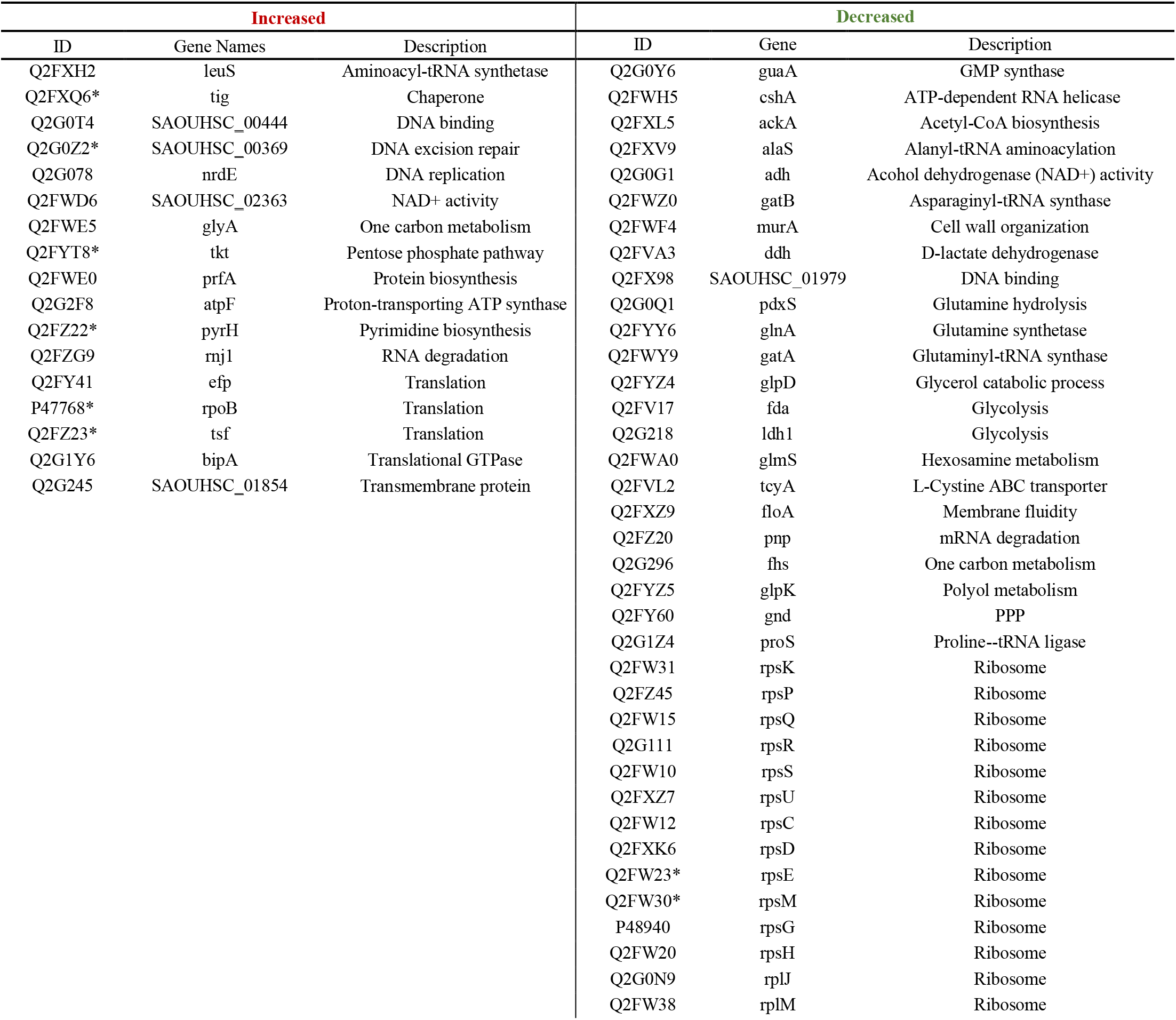
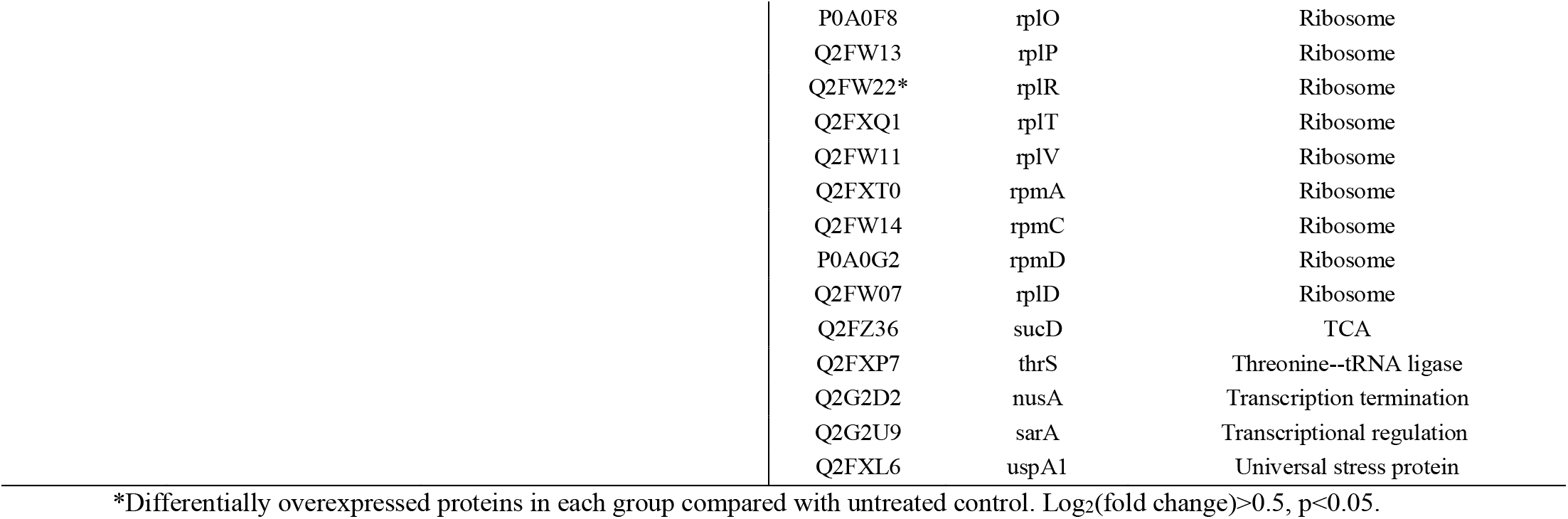
Increased and decreased proteins in enrofloxacin persisters.

**Figure 3.**
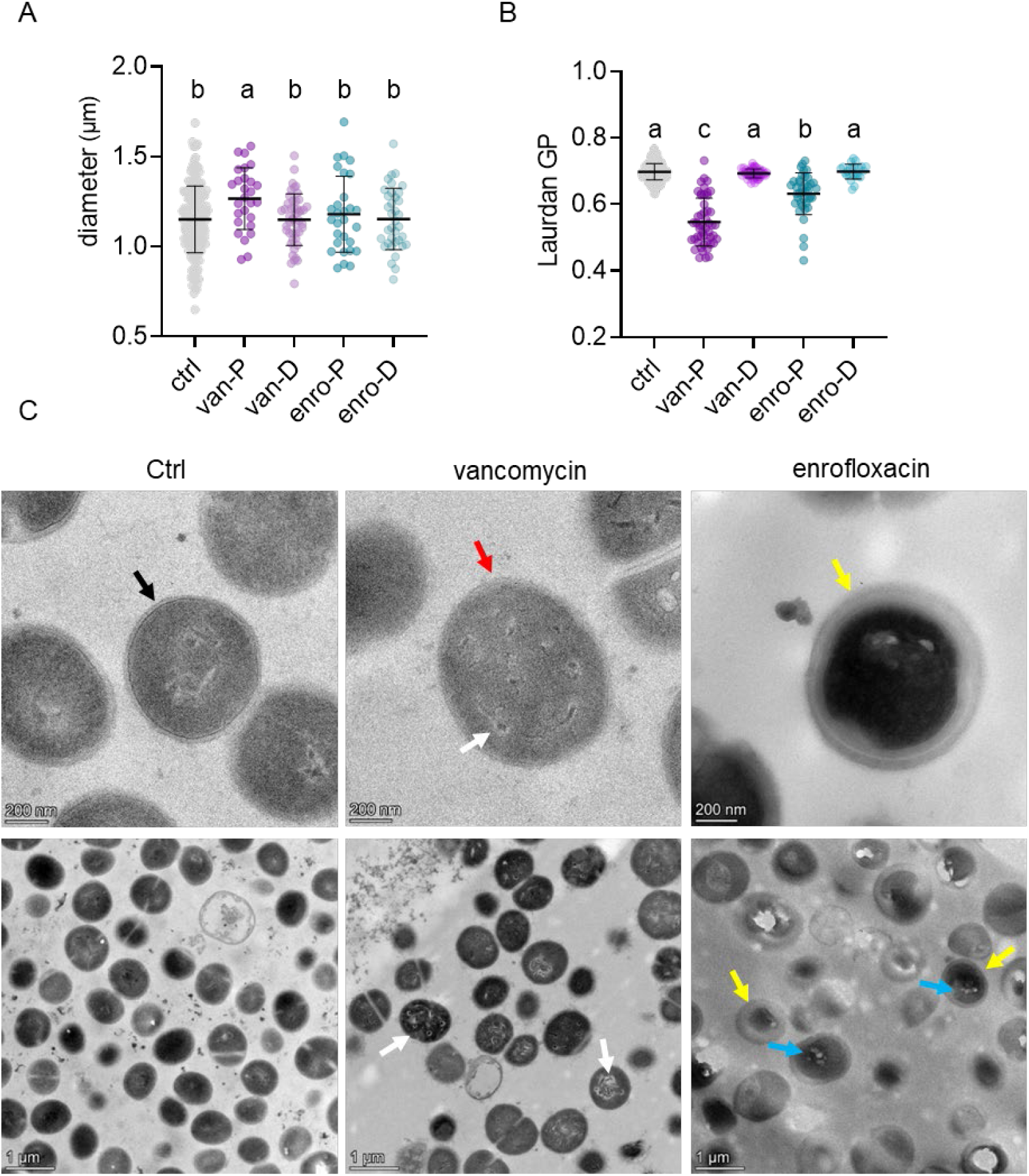
Alteration in morphology before and after antibiotic exposure. **A**. Diameter of vancomycin induced persisters (van-P), vancomycin killed cells (van-D), ctrl, enrofloxacin induced persisters (enro-P) and enrofloxacin killed cells (enro-D). Vancomycin induced persisters are slightly bigger than control cells, dead cells and enrofloxacin induced persisters. **B**. Quantified Laurdan GP shows a more fluidified cell membrane in persisters than in vegetative cells. Besides, vancomycin persisters have more fluid membranes than enrofloxacin persisters. **C**. Transmission electron microscopic images of vegetative *S. aureus* cells (ctrl), and vancomycin and enrofloxacin treated cells. Black arrow: normal cell structure. Red arrow: vancomycin treated persisters without tripartite-layer cell wall. White arrows: protein aggregations. Yellow arrow: thickened cell wall. Blue arrow: mesosomes-liked structures.

In both van-P and enro-P, increased proteins are mainly associated with ATP binding, translation, central carbon metabolism, self-digestion, anti-oxidation and chaperones, while some decreased proteins involved in ATP-binding, translation, and central carbon metabolism were also observed. The overexpression of many ATP-binding proteins illustrates the energy requirement of *S. aureus* persisters to combat antibiotic exposure. In van-P, we detected 14 increased ATP binding proteins including 9 translation related proteins, 3 chaperones and 2 transmembrane transporters. In contrast, three decreased ATP-binding proteins were identified including GuaA and gatB that both related to glutamine consumption. This could be due to increased peptidoglycan biosynthesis where glutamine is an essential precursor [23]. In enro-P, 4 increased ATP-binding proteins were Q2FZ22 and Q2G078 that are associated with DNA replication and repair, Leucyl-tRNA synthetase Q2FXH2 as well as proton-transporting ATP synthase Q2G2F8. These data suggest that ATP in persisters was mainly used for protein synthesis and modulation as well as transmembrane transport. In enro-P, DNA repair also played an important role to survive DNA damaging enrofloxacin, which will be discussed further later on.

Meanwhile, distinctive predominant proteins related to protein synthesis were identified with either increased or decreased expression levels. Interestingly, increased proteins that contribute to protein synthesis are mainly involved in aminoacylation (asnS, aspS, gltX, leuS, lysS or serS in van-P; leuS in enro-P), transcription elongation (efp and nusG in van-P; tsf and efp in enro-P) and ribosome assembly (bipA and engA in van-P, bipA in enro-P), while decreased translation related proteins are mostly ribosomal proteins (20 ribosomes in van-P and 23 in enro-P). This indicates that while general protein synthesis is inhibited in persisters, the modulation of specific protein is still actively happening. The elevated aminoacylation, such as mediated by SerS, GltX and LysS, could also contribute to peptidoglycan biosynthesis and/or membrane phospholipid modification [24,25].

Central carbon metabolism plays a crucial role in providing an energy source and precursors. In van-P, we identified 8 increased proteins participating in glycolysis (pgi), in carbon (1C) metabolism (glyA), in the pentose phosphate pathway (PPP) (zwf, cidC) and in the TCA cycle (citB, citZ, sucC, citC), as well as 2 decreased proteins involved in the conversion of lactate and pyruvate (ldh1 and ddh). In enro-P, only GlyA and Tkt in the PPP were increased while SucD (TCA), Fhs (OCM), Gnd (PPP), Fda, Ldh1 and Ddh (glycolysis) proteins were decreased. These results suggest differential carbon metabolism activities between these two conditions. Interestingly, neither of the proteins increased or decreased in expression function as ATP generator, suggesting that the ATP generation ability is likely similar as in vegetative cells at the enzyme level, but the process may be repressed due to factors such as generally decreased metabolic activity, lack of reaction substrates, as well as possible reorientation of the respiratory chain caused by anti-oxidative stress response as discussed later.

Self-digestion or autophagy of cells of a subpopulation, including the intracellular degradation of RNA, protein and lipids, is an important survival strategy for a bacterial population to combat unfavored conditions through recycling of energy harvesting molecules or acceleration of dormancy [26]. In *E. coli* stationary phase populations, a correlation between self-digestion and persister formation was proposed [27]. In our work, we used an early log phase population to generate persisters where starvation is unlikely to occur. However, we identified the overexpression of self-digestion related proteins, including proteases glutamyl aminopeptidase, HtrA1 and ClpP in van-P, as well as RNase J1 in both van-P and enro-P persister samples. Proteases are of great importance in protein homeostasis and antitoxin degradation, and in this way contribute to persister formation [28]. On the other hand, the overactivation of proteases by acyldepsipeptide antibiotics like ADEP4, was found to cause dysregulation of proteolysis, excessive self-digestion and eventually persister death [29,30]. RNase J1 is a 5-3′ ribonuclease in *S. aureus* that is associated with RNA decay and maturation, as well as exoribonucleolytic activity [31]. The contribution of RNA degradation to persister formation has been related to starvation [32] or the toxin-antitoxin system [33]. The exact relationship between RNase J1 and *S. aureus* persisters has not been fully assessed yet, but we hypothesize that RNnase J1 may contribute to persister formation by degrading unnecessary RNAs, thereby decreasing the general metabolic activity. The activation of self-digestion proteins and stated increased translation-related proteins underscores the importance of maintaining a delicate metabolic balance in persister formation.

Antioxidative proteins were observed in both persister populations, which indicates increased levels of reactive oxygen species (ROS) during vancomycin and enrofloxacin exposure. Specifically, 9 proteins were found that contribute to oxidative stress response in van-P, including H_2_O_2_ sensitive Q2FWD6 and KatA [34], disulfide bond control protein AhpC [35] and TrxB [36], Fe-S cluster assembly protein Q2FZY3 and CysM [37], OSR regulator YchF [38], as well as NADPH generating proteins CitC and G6PD. Besides, the decrease of lactate dehydrogenase in both persistent populations indicated that they were not under nitric oxide (NO·) stress [39]. In comparison, of these only protein Q2FWD6 was significantly increased in enro-P. The overexpression of oxidative stress relevant proteins suggests that persisters requires them to combat high level of several ROS caused by the bactericidal antibiotic vancomycin and enrofloxacin.

In response to the cell wall targeting antibiotic vancomycin, *S. aureus* persisters increased cell wall biosynthesis related proteins, including Pbp2 and SerS that contributed to peptidoglycan biosynthesis [24], as well as Tarl facilitating teichoic acid biosynthesis. In addition to stated proteins that directly stimulate cell wall biosynthesis, aspartate aminotransferase AspB, chaperone PrsA and flotillin protein FloA, demonstrated to be indirectly associated with peptidoglycan synthesis [40–42], were also increased. Amongst the primary functions of FloA is mediation of membrane fluidity which may also indicate a distinct characteristic of van-P.

In enro-P, we observed the stimulation of proteins related to DNA repair and replication. Precisely, protein Q2G0Z2 contributes to DNA excision repair. Q2G0T4 is associated with the RecF pathway which displays homologous recombination to repair DNA. Additionally, Q2G078 and Q2FZ22 provide essential precursors for DNA synthesis. Likely through elevated DNA replication and repair, enrofloxacin persisters were able to survive enrofloxacin treatment and maintain the ability to be resuscitated after stress removal.

### Morphological characterization

To demonstrate the consequences of proteome profile alteration obtained from isolated persisters, we conducted various tests at both single-cell and population level.

Proteomic analysis revealed several proteins that are related to morphological changes such as peptidoglycan biosynthesis in van-P that might cause thicker cell wall. Additionally, proteins associated with membrane composition and transmembrane transport were identified in both persister populations, suggesting potential interference in membrane features, including membrane fluidity. Therefore, phenotypic characteristics of persisters were analyzed microscopically.

With CFDA-PI double staining, persisters and control vegetative cells were labeled with only green fluorescence while dead cells displayed red color. Single cells were first observed using fluorescence microscopy. The measurement of cell diameter (Figure 3A) showed that van-P cells were slightly bigger (average Δ diameter ≈ 0.13 μm) than vegetative cells and enro-P, while no detectable differences were observed between truly ‘killed’ cells and control cells. We also measured cell membrane fluidity using a polarity sensitive fluorescent probe, Laurdan (6-Dodecanoyl-2-Dimethylaminonaphthalene). Combining with CFDA and PI, the membrane fluidity of single persistent, dead and control cells was determined by the Laurdan generalized polarization (GP) value (Figure 3B) where a higher value represents lower fluidity. Consistently, we found that while cells killed by either vancomycin or enrofloxacin showed no fluidity change compared to control cells, persisters on average had a more fluid membrane, especially the van-P.

In addition, we performed transmission electron microscopy (TEM) of untreated control cells and of 24h vancomycin- and enrofloxacin-treated samples that contained 99.1 %, 86.9% and 77.9 % of CFDA^+^-PI^-^ cells (analyzed through flow cytometry before fixation), respectively. Because of the high proportion of persisters, intact cells in the antibiotic treated samples in these images were considered to be persisters. Distinctive morphological features were observed in each group (Figure 3C). Vancomycin treated cells displayed an altered cell wall without the typical tripartite-layer structure (red arrow), in contrast to control cells that had a normal cell structure (black arrow). Additionally, inclusion bodies structures and areas [59], were observed in the majority of intact cells (white arrows). This suggests that protein aggregation was likely associated with the formation of these vancomycin persisters. This result aligns with our previous observations that vancomycin persisters showed a high level of protein aggregation [8] and provides a potential explanation for the upregulation of chaperones which are associated with protein disaggregation. However, there was no noticeable increase in cell wall thickness despite the observed upregulation of proteins associated with cell wall synthesis in the proteomics data. Interestingly, enrofloxacin treated cells prominently exhibited thickened cell walls (yellow arrows). This suggests that increased cell wall thickness is a general stress response of *S. aureus* occurring during enrofloxacin exposure, not necessarily linked to persisters, and may explain why no cell wall synthesis proteins are found to be increased in the enro-P proteomic data. Additionally, mesosomes-liked structures (blue arrows) were observed in enrofloxacin treated cells which are associated with antibiotic survival and membrane alterations [60,61].

The detected alterations in cell size, membrane fluidity, protein aggregation and cell morphology of the persisters, which were neither observed in vegetative nor in dead cells, demonstrate that these alterations were not only caused by antibiotic treatment. Instead, they were attributed to the stress responses of persisters to survive given stresses, especially vancomycin, which potentially contribute to persisters formation and maintenance.

### Metabolic characterization

Contrary to the general contention that persister cells are metabolically inactive, we demonstrated in previous work that glucose consumption (Figure 4A) and/or acetate generation (Figure 4B) still gradually occurred within stable van-P and enro-P populations, with van-P displaying higher activity in consuming glucose than enro-P [8]. Based on these observations, we conducted further investigations to assess the metabolic alteration of persisters by examining biochemical changes in the culture medium and within FACS-isolated cells.

**Figure 4.**
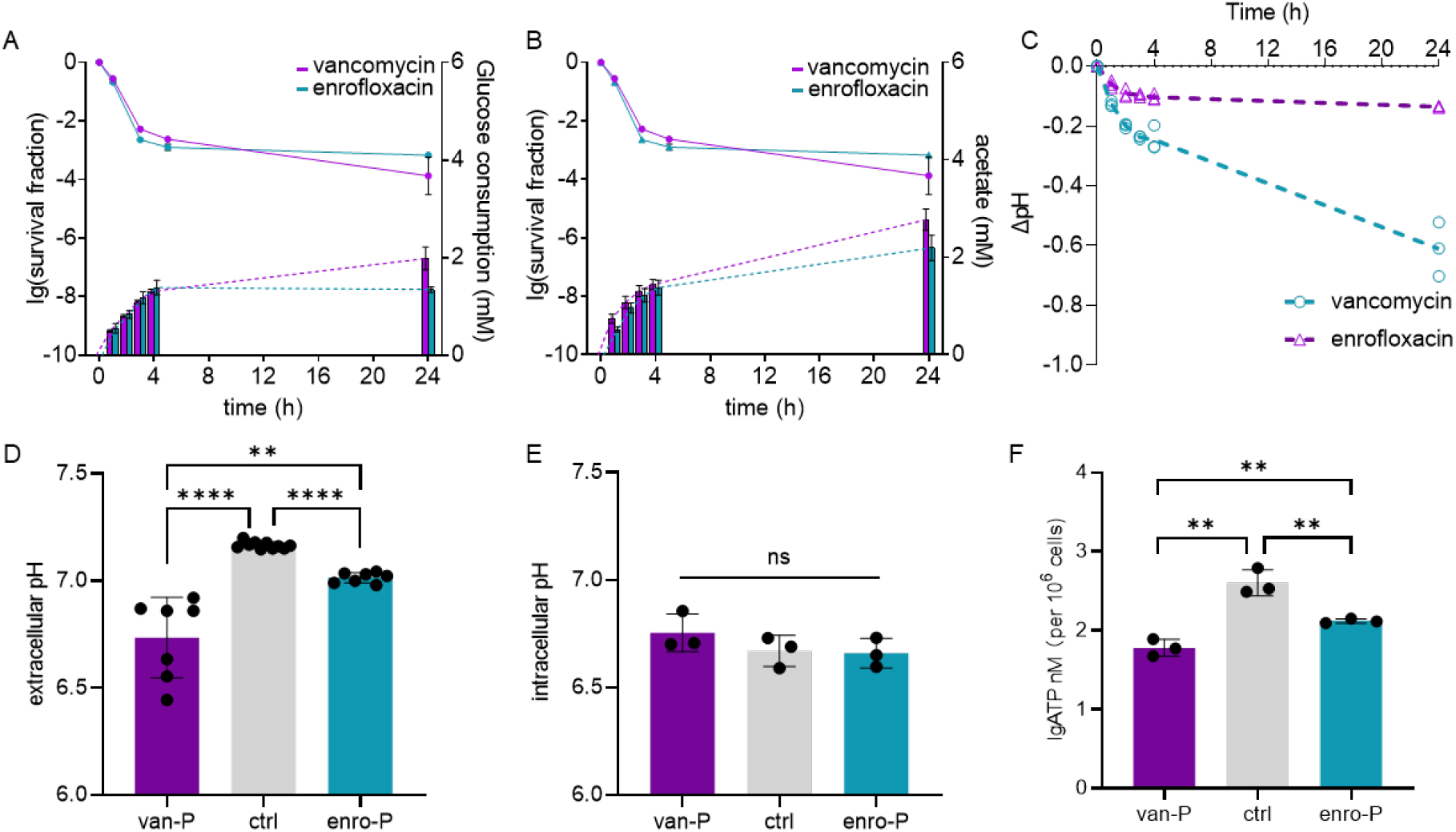
Alteration in metabolic activities of persisters detected both extracellularly and intracellularly. Extracellularly, we tested the glucose consumption (**A**), acetate generation (**B**) and pH alteration (**C**) during persister formation. The comparison between extracellular pH before and after 24 hours of antibiotic exposures were shown in (**D**). With 10^6^ FACS isolated cells in each treatment, we tested the difference in intracellular pH (**E**) and intracellular ATP level (**F**) between persisters and vegetative cells. Noted figures A and B are cited from our previous work [8].

Firstly, we observed a pH drop in the medium of all antibiotic treated samples (Figure 4C). Notably, vancomycin treatment led to a more pronounced decrease in pH even during the stable persister phase while enrofloxacin caused a pH drop mostly within 4 hours of exposure. In 24h treated samples, vancomycin treatment resulted in a more acidic environment than in the initial culture while enrofloxacin treatment caused a significant but only minimal pH reduction (Figure 4D), which could be due to the release of acetate during carbon metabolism. The medium pH did not affect the intracellular pH as no significant pH differences were observed between isolated persisters and control cells (Figure 4E). Subsequently, the active carbon metabolism and the overexpression of ATP consuming proteins observed in the proteomics data prompted us to examine intracellular ATP levels. Reduced intracellular ATP levels have been confirmed as a crucial contributor to persister formation in various bacterial species including *S. aureus* [62,63]. Confirming our expectations, our persisters contained lower ATP levels compared to vegetative cells (figure 4F). Interestingly, van-P contained less ATP than enro-P. Considering the higher glucose consumption and this lower level of intracellular ATP observed in van-P, we propose that the vancomycin treated cells likely have higher metabolic activity than enrofloxacin treated cells. This may be the reason of faster resuscitation of van-P as shown in Figure 1C. Additionally, the observation of more active ATP generation but lower intracellular ATP levels in vancomycin persisters indicates that it may require more energy to generate vancomycin persisters, which also matches our findings in proteomics that van-P express more ATP-consuming proteins.

The effect of ATP repression, protein synthesis inhibition and ROS levels on persister formation.

Through the proteomics analysis, we identified distinctive predominant proteins that were stimulated or repressed in van-P and enro-P. Within these, both vancomycin and enrofloxacin persisters have more ATP-dependent proteins and decreased translation related proteins. Therefore, we assumed that *S. aureus* persisters have a noticeable level of energy requirement to form persisters. Additionally, the inhibition of translation might lead to higher level of persisters. To test this assumption, we used *S. aureus* USMA-1, together with two other clinically-isolated *S. aureus* strains JAR 06.01.31 and 42D, to generate persisters upon exposure to 100-fold MIC of carbonyl cyanide m-chlorophenyl hydrazone (CCCP). This compound fully eliminates ATP synthesis by dissipating the proton motive force. We also deployed the antibiotic tetracycline which prevents the binding of aminoacyl-tRNA to the 30S ribosome, blocking translation but does not affect ATP synthesis. Time kill curves show that under high levels of CCCP exposure (Figure 5A), only few cells in the USMA-1 population formed colonies, and no detectable alive cells were found with the other two *S. aureus* strains. This demonstrates that high levels of CCCP which potentially completely eliminated ATP production, prevent *S. aureus* persister formation. This suggests the significance of maintaining a certain level of cellular ATP for persister formation.

**Figure 5.**
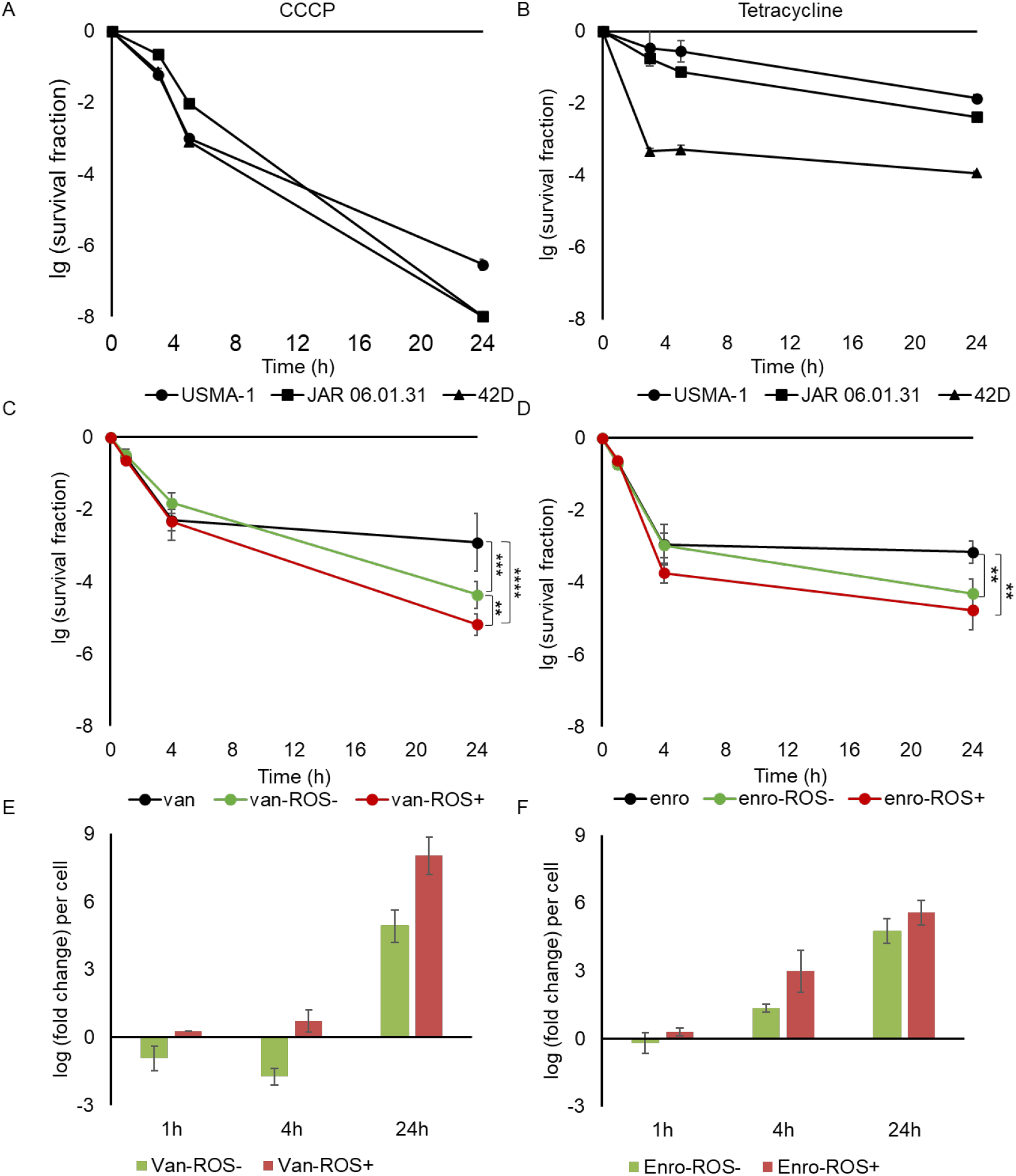
Time kill curves of *Staphylococcus aureus* populations exposed to different stressors. To analyze the importance of ATP production and protein synthesis in persister formation, 100-fold MIC of CCCP (**A**) or tetracycline (**B**) was used to treat three *S. aureus* strains USMA-1, JAR 06.01.31 and 42D. To test the influence of ROS levels on persister formation under vancomycin and enrofloxacin exposure, time kill assay was performed with USMA-1 populations exposed to thiourea (reducing ROS; ROS-) or H_2_O_2_ (increasing ROS; ROS+) before the treatment with vancomycin (**C**) or enrofloxacin (**D**). The ROS level of each sample was also monitored before and after 1h, 4h and 24h of vancomycin (**E**) or enrofloxacin (**F**) exposure using luminescent probe L-012. The data were normalized based on the number of intact cells (including persisters, VBNC cells and dead cells) and presented as log (fold change) compared with the ROS level of “antibiotic only” cultures at each time point. Minimum detectable survival fraction is 10^-8^. n≥3. **: p<0.01. ***: p<0.001. ****: p<0.0001. n=3.

Interestingly, when treated with high levels of tetracycline (Figure 5B), the antibiotic biphasic kill curve, the hallmark of persister formation, was only observed in the 42D population, resulting in ∼0.01 % surviving cells which was about 10-fold less than van-P and enro-P. Oppositely, slow but consistent killing was observed in USMA-1 and JAR 06.01.31 samples, resulting in a more antibiotic tolerant population with a higher survival fraction after 24 hours of treatment. This result shows that translation block by tetracycline did not cause higher persister levels as no biphasic killing curve was seen, but likely did contribute to antibiotic tolerance and thereby to increased cell survival.

Both glycopeptides and quinolones have been found to induce the accumulation of the hydroxyl radical (·OH) in *S. aureus* [64]. Different types of oxidative stress related proteins were found overexpressed in van-P, which indicates that the ROS level might affect persister generation, especially under vancomycin stress. Therefore, we performed a time kill assay to study the effect of ROS levels on persister formation under vancomycin and enrofloxacin treatment. Through pretreating the culture with 150 mM of the ROS scavenger thiourea [65] or with 50 μM of the ROS precursor H_2_O_2_ [66] before antibiotics exposure, we created ROS depleted (ROS-) and ROS increased (ROS+) environments, respectively, for each sample. Interestingly, compared with control samples that had no ROS modulation, both ROS- and ROS+ conditions resulted in less survival after 24h of vancomycin (Figure 5C) or enrofloxacin (Figure 5D) treatment, with vancomycin exposure causing larger differences. Additionally, increased intracellular ROS levels cause more killing in vancomycin than in enrofloxacin treated samples. We also monitored the dynamics of ROS levels before and after 1h, 4h and 24h of vancomycin (Figure 5E) and enrofloxacin (Figure 5F) treatment using L-012, a luminol-based luminescent probe for ROS detection [65]. The ROS level of each sample was quantified and normalized to the ROS level of “antibiotic only” cultures at each time point. As expected, the ROS level per cell in ROS+ samples quickly increased while ROS-cells displayed a decreased ROS level compared with the “antibiotic only” samples at the beginning of antibiotic exposure, especially in vancomycin samples. In 4h enrofloxacin treated, and the 4 and 24h vancomycin treated samples, higher ROS level were detected in ROS-samples which could be due to the complete consumption of thiourea. Interestingly, after 24h of incubation, cells in both ROS- and ROS+ samples contained more ROS than control “antibiotic only” groups samples. This is likely linked to the killing in these two groups compared with control cells. In enrofloxacin treated samples, induced ROS also likely initiated the SOS response that then stimulates DNA repair proteins which were found highly expressed in enro-P.

## Discussion

In this work, we validated the persister isolation method in antibiotics treated *S. aureus* populations. With this effective persister isolation method, we separated persister populations from other potentially existing cell types in antibiotic treated samples, including debris, dead cells and VBNC cells for further analysis. The results enabled us to conduct in-depth molecular and phenotypical analysis specifically on persisters.

### Proteomics application in persister-research

With the proposed persister isolation method we previously described for *Bacillus subtilis* and with minor modification applied for *S. aureus* in the present work, pure persister populations could be enriched through simple double staining and FACS separation. Nonetheless, one limitation of the present proteomics work is that FACS flow rate and the need to minimize sorting time to prevent sample damage resulted in small sample size. This consequently caused a relatively low proteome sequence coverage, potentially restricting the detection to only highly induced or repressed proteins. Therefore, to obtain the maximum utilization of limited protein detection, we not only analyzed the uniquely expressed proteins in persistent populations and considered them as persister-related proteins, but also discussed those proteins that were detected in control cells but missing in persisters, which we thus considered repressed proteins. The high similarity between three biological replicates demonstrated that the lower proteome coverage (protein sequence depth) did not compromise the quality of the proteomics data, demonstrating that the presented proteomics results accurately represented the main changes occurring in tested persisters.

Compared to previous proteomics results, such as one that identified 710 proteins in rifampin and ampicillin treated *Escherichia coli* enriched population by removing dead cells through magnetic-beads-based separation [11], or another one analyzing 903 proteins obtained directly from *Candida albicans* biofilm samples, our finding with 451 detected protein appear relatively modest. However, considering the specific research objectives of describing the dynamics of proteome features observed during persister formation and in isolated persisters, well separated from VBNC cells, our findings provide more precise information. Moreover, the isolation method we employed enables the application of single-cell sequencing techniques, which could further enhance the precision of persister-related research.

### Morphological alteration in persisters

When investigating morphological characteristics with both vancomycin and enrofloxacin persisters, we observed significant size change in van-P, and membrane fluidity change in both persister populations, while no detectable changes were found in killed cells. This demonstrated that the alterations were related to not only antibiotic exposure but more to stress responses that led to persister formation, facilitating *S. aureus* persister survival. Interestingly, the observed enlargement of vancomycin persisters was not attributable to a thickened cell wall as expected. As shown by TEM images, vancomycin persisters have a changed cell wall without the typical tripartite-layer structure. On the other hand, enrofloxacin triggered a thickened cell wall in both intact and compromised cells, suggesting that increasing cell wall thickness may serve as a common defensive mechanism against enrofloxacin induced antibiotic stress.

Membrane fluidity change has been reported as an adaptive stress response against environmental stressors including temperature, pressure stress, as well as antimicrobial agents [67,68]. We observed that particularly vancomycin generated persisters have a more fluid membrane. In van-P, we found the overexpression of FloA, which likely contributes to increased fluidity [40]. In agreement with these observations, membrane lipid analyses indicated that 8-24h of incubation with high level vancomycin resulted in more fluid membranes in *S. aureus* [69]. However, in enro-P that show a slightly more fluid membrane than untreated cells, the expression of FloA was decreased. This implies that membrane fluidity alteration might also be attributed to non-lipid alterations such as the overexpression of membrane ATPase and other membrane proteins, including various metabolite transporters or enzymes. Further investigations into the underlying mechanisms regulating membrane fluidity and its impact on persister physiology are needed for a better understanding of persister formation and survival strategies. Besides, membrane composition alteration likely also results in change of other membrane features such as electric potential, permeability and net charge, which could potentially interfere with the sensitivity of persisters towards certain antimicrobial agents e.g., cationic antimicrobial peptides [70].

### The balance in persister formation

This research has emphasized the importance of maintaining a delicate balance in persister formation. One factor is an optimal intracellular ATP level. The overexpressed levels of ATP-dependent proteins, the presence of active but slower carbon metabolism, and the block of persister formation by CCCP exposure all indicate that *S. aureus* cells required a certain amount of energy to form persisters, particularly when exposed to vancomycin stress. Consistent with our findings, many studies have observed that decreased ATP is a common characteristic and contributor to persister formation in various bacterial species [63]. However, excessive or even complete depletion of ATP can either prevent persister formation as we showed, or can lead to an excessive level of protein aggregation that causes cell death or deeper dormancy [71].

Another aspect is the balance of translation. Proteomics data shows that both persister samples contained less ribosome proteins, but increased levels of several proteins related to aminoacylation and elongation, together with the overexpression of self-digesting proteins. These results suggested that repressed general translation helps to form growth arrested persisters. Meanwhile, specific translation and its regulation still occurred during persister formation. This highlights the importance of maintaining a balanced translation process. When treated with tetracycline that directly inhibits translation, *S. aureus* USMA-1 and JAR 06.01.31 strains show relatively slow but consistent killing with more survival after 24h of incubation, forming a tolerant-like population, suggesting that the blocked translation quickly drove cells into dormancy, but at the same time, failed to trigger persister formation. In 42D populations, however, tetracycline did trigger the generation of persisters. This difference in persister formation could be due to the genetic differences between the strains. This observation reminds us that persister formation is a multi-factorial process, dictated by the nature of the stressors as well as influenced by the genetic and regulatory differences between bacterial strains which may lead to strain-specific stress response against one stressor.

The role of ROS in antibiotic-mediated killing and the effect of ROS alteration in bacterial survival have been intensively researched but are still controversial and are highly depended on the experimental conditions, as discussed in [72]. In the present work, the alteration of ROS levels and corresponding surviving fraction demonstrated the importance of ROS balance in persister formation. This was demonstrated by the findings that both vancomycin and enrofloxacin treated samples displayed consistent levels of persisters at 4h and 24h of treatment which represents the period of stable persistent populations, and both artificially increasing and decreasing ROS level resulted in less cell survival and continued cell killing over time. Besides, when normalized to the ROS level per intact cell (including persisters, VBNC and dead cells), higher ROS levels were observed in even ROS-samples after 24h incubation than “antibiotic only” treated cells. This indicates that when cells did not initiate the oxidative stress at the start of the ROS-triggering antibiotics exposure due to the addition of thiourea, the persister inducing regulating system was then interrupted during prolonged exposure. Conversely, within host cells, higher ROS levels have been correlated with increased *S. aureus* oxacillin persister generation [21]. Additionally, the increase of intracellular ROS level by menadione, which mimicked the phagocyte-induced ROS production, resulted in an almost completely tolerant *S. aureus* population to rifampicin [65]. These contrasting outcomes can be attributed to the different kinds and levels of oxidative stress bacteria encounter and the distinct modes of action of tested antibiotics, highlighting the necessity of considering specific experimental settings when dealing with persisters through ROS regulation.

In conclusion, this work proved the efficiency of our proposed method for isolation of persister from antibiotic treated *S. aureus* populations, making the studies on the molecular physiology of pure *S. aureus* persister populations feasible. Thus, further experiments such as using mutated strains to validate certain proteins or metabolic pathways of interest will be possible owing to our methodology.

The characterization experiments highlighted unique features of specific persister populations and the importance of ensuring a balanced stress response capable of persister formation, which could provide valuable information in persister prevention or eradication of potential clinical use.

## Methods

### Bacterial strains

Three clinically isolated *S. aureus* strains were used in this work. Specifically, *S. aureus* ATCC49230 (UASM-1) has high virulence, initially isolated from a patient with chronic osteomyelitis [73]. *S. aureus* JAR 06.01.31 was isolated from an infected human hip prosthesis [74]. *S. aureus* 42D is also a clinically isolated strain with relatively low-virulence [75]. All strains were incubated in Tryptic soy broth (TSB) medium at 37 ℃ with constant rotation at 200 rpm when needed or as otherwise specified. Single colonies of three strains were inoculated from TSB agar plates into liquid medium for overnight incubation. Afterwards, overnight cultures were 1:100 transferred into fresh medium and incubated until early log phase (OD_600_=0.4-0.6) for the following experiments. For antibiotic exposure, cultures were diluted with TSB to OD_600_=0.2 before antibiotic treatment.

### Minimal inhibitory concentration (MIC) measurement

Four antimicrobial agents (all Sigma-Aldrich, Zwijndrecht, the Netherlands), vancomycin, enrofloxacin, CCCP and tetracycline were used to generate persisters. MIC of every antimicrobial compound was determined. Specifically, 150 μl mixtures including around 10^6^ *S. aureus* cells from early log phase culture and two-fold serial dilutions (from 128 μg/ml to 0.125 μg/ml) of antimicrobial compounds were added to each well of the 96 well plate. Control groups contained only *S. aureus* culture and negative control samples included only TSB medium). During 18h incubations, OD_600_ measurement in a microplate reader (Multiskan™ FC, Thermo Scientific, Etten-Leur, the Netherlands) where performed. The minimal concentration that caused non-measurable OD_600_ change was determined as MIC. Three biological replicates were performed, and the results were listed in table 3. MIC measurement was also used to recheck the absence of resistant *S. aureus* after antimicrobial exposure.

**Table 3.**
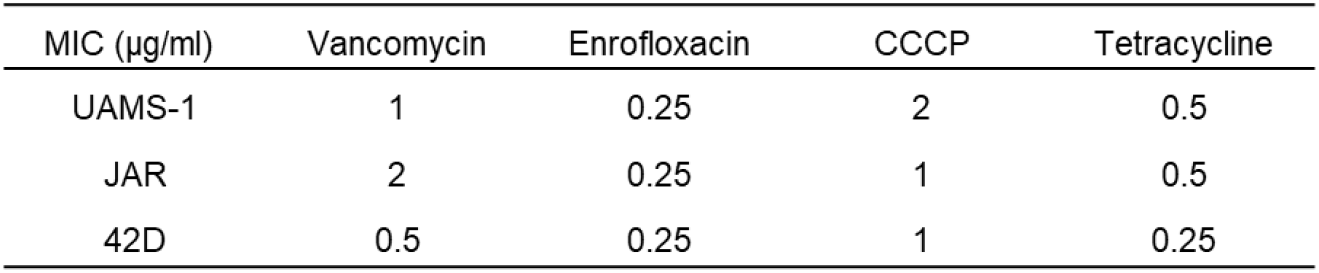
MICs for each strain for each of the four antimicrobial agents.

### Time kill assay

Time kill assay was performed by exposing early log phase cultures of each strain to 100-fold of the corresponding MIC of every antimicrobial agent. Before and after a certain time of incubation, 1 ml samples were transferred to 1.5 ml Eppendorf tubes and were then centrifuged at 4000*g for 5 mins and washed twice by either isotonic salt solution or Phosphate-buffered saline (PBS) to remove remaining antibiotics. The number of surviving cells were quantified by 10-fold dilution and drop plating. Time-kill curves were made for each treatment.

### Double staining and persister isolation

We tested previously proposed persister isolation method [10] with selected persistent populations. Similarly, after CFDA and PI staining, antibiotic treated samples were either inoculated into either TSB or isotonic salt solution as negative control. The CFDA and PI signals were then detected with 488 nm excitation and 519 nm emission, or with 532 nm excitation and 617 nm emission, respectively.

Unstained samples containing 1% dimethyl sulfoxide (DMSO, Sigma-Aldrich, St Louis, USA) were used to set up the gates. Later on, persisters were detected as CFDA^+^-PI^-^ by flow cytometry (FACSAria™ III, BD Biosciences, San Jose, CA) and microscopes. FACS were used for isolation when needed. Untreated early log phase culture was used as control in following experiments.

### Proteomics sample preparation and LC-MS analysis

Persistent populations generated by 24h of vancomycin and enrofloxacin treatment were washed twice with PBS and double stained for FACS isolation. For each sample, around 3*10^^7^ persisters were collected. Afterwards, bacterial cells were concentrated and resuspend in 1% SDS. Protein extraction, sample preparation, LC-MS measurement and data analysis were performed following the previously described protocol [8].

### Microscopy

The morphological features of persisters were performed with transmission electron microscope (TEM). Specifically, 15OD_600_ culture before and after 24h vancomycin or enrofloxacin exposure were collected and washed twice. The proportion of CFDA^+^-PI^-^ cells in each sample were detected by flow cytometry before fixation. The pallet of each sample was then fixed in Mc Dowell containing paraformaldehyde and glutaraldehyde (Polysciences, Inc., Warrington, PA, USA) and post-fixed with 1% osmium tetroxide (OsO4, Electron microscopy sciences, Hatfield, PA, USA). Subsequently, the samples were dehydrated in an alcohol series and embedded into Epon (LX-112 resin Ladd research, Williston, VT, USA). Ultrathin (60 nm) epon sections of the samples were cut and collected on Formvar-coated grids, counterstained with uranyl acetate and lead citrate. Sections were examined with a Talos L120C electron microscope (Thermo Fisher Scientific, Eindhoven, The Netherlands), images were taken with a BM-CETA camera using Velox software (Thermo Fisher Scientific, Eindhoven, The Netherlands). Duplicates were used for TEM.

Cell size, membrane fluidity and intracellular pH were conducted with Eclipse Ti microscope (Nikon, Tokyo, Japan) equipped with phase-contrast and fluorescence components. For membrane fluidity, double-stained cells were also stained with 10 μM Laurdan (6-dodecanoyl-N,N-dimethyl-2-naphthylamine, Sigma Aldrich, Zwijndrecht, the Netherland). Laurdan was excited at 395 nm and 470 nm emission wavelength was used for rigid membrane and 508 nm for fluid membrane environment. Laurdan GP of the microscopic images and its quantification were performed using the CalculateGP ImageJ plugin (https://sils.fnwi.uva.nl/bcb/objectj/examples/CalculateGP/MD/gp.html). Intracellular pH detection was performed based on [76] with the modification in using PBS for washing and pH calibration curve.

### ATP level, pH and ROS measurement

For both intracellular ATP level and pH, CFDA^+^PI^-^ cells were isolated by FACS and 10^6^ cells were used for one test. The detection of intracellular ATP level was conducted by BacTiter-Glo™ Microbial Cell Viability Assay (Promega Corporation, Leiden, the Netherlands). ATP disodium salt (Sigma-Aldrich, Zwijndrecht, the Netherlands) was dissolve in sterile water and ten-fold serially diluted (1nM to 10μM) to generate standard curve. Luminescence values were detected by a BioTek SynergyMx microplate reader. Intracellular pH was performed following the protocol of Nielson *et al*. [76] with the modification of using PBS as reaction solution in all samples and measure the intensity with the BioTek SynergyMx microplate reader. Extracellular pH was measured with a Lab pH meter inoLab® pH 7110 at each timepoint with filtrated medium. To create ROS- or ROS+ environment to analyse the impact of ROS levels with respect to persister formation, thiourea (VWR International B.V., Amsterdam, the Netherland) or H_2_O_2_ (Sigma Aldrich, Zwijndrecht, the Netherland) were added in the culture 20 minutes before the addition of antibiotics, with the final concentration of 150 mM or 50 μM, respectively. ROS level was measured with L-012 as described in [65] and measured with the BioTek SynergyMx microplate reader. Then the luminesce units were used to present ROS level by normalizing with the number of all intact cells at each corresponding time point and visualizing as log_2_ (fold change) compared with the ROS level of “antibiotic only” cultures at each time point.

## Data availability

All mass spectral data related to this publication have been deposited in the MassIVE repository (https://massive.ucsd.edu/ProteoSAFe/static/massive.jsp) under the dataset identifier MSVXXXX

## Author Contributions

Conceptualization, S.B., S.A.J.Z. and S.L.; performing experiments and analyzing data, S.L., P.L., S.J., N.W, G.K.; writing—original draft preparation, S.L.; writing—review and editing,

G.K. S.B. and S.A.J.Z. All authors have read and agreed to the published version of the manuscript.

## Acknowledgments

We acknowledge Gonzalo Congrains Sotomayor for his expert technical assistance in flow cytometry experiments. S.L. acknowledges the China Scholarship Council for her PhD scholarship.

